# Dpp and Immune Response Pathways Factors Mediate Paracrine Induction of Senescent Cells in *Drosophila*

**DOI:** 10.64898/2025.12.11.693679

**Authors:** Juan Manuel Garcia-Arias, Mireya Ruiz-Losada, Natalia Azpiazu, Ginés Morata

## Abstract

Transition toward senescence is a cellular response to different stressors like ionizing radiation, telomere shortening or oncogene activation. This phenomenon is evolutionarily conserved across species, from insects to humans. Senescent cells (SCs) permanently withdraw from the cell cycle and undergo a series of physiological changes, most notably the acquisition of a robust secretory activity characterized by the release of numerous molecules, including cytokines, chemokines, and metalloproteinases. Through this program, termed Senescence-Associated Secretory Phenotype (SASP), SCs actively communicate with and influence their microenvironment. In mammalian tissues the number of SCs increases with age and their accumulation has been proposed to contribute to several age-associated pathologies. Studies in vertebrate systems have demonstrated that new SCs can arise through paracrine signaling from pre-existing SCs, a process that requires the activity of Transforming Growth Factor β (TGF-β).

We have investigated the phenomenon of paracrine recruitment of SCs in the *Drosophila* wing disc. Our results show that an initial stress event induces a primary wave of SCs, comprising approximately 10% of the target cell population. Subsequently, a second wave of SCs emerges through paracrine signaling from the initial cohort, increasing the overall proportion of SCs to about 24%. The formation of this second wave is mediated by the growth factor Decapentaplegic (Dpp), the *Drosophila* ortholog of TGF-β. Dpp activates a non-canonical signaling route in non-SCs, driving their conversion to a senescent state. This novel branch of the Dpp pathway engages several components of the innate immune response.

Collectively, these findings underscore the evolutionary conservation of senescence-associated signaling networks and suggest that paracrine amplification of senescence may play a role in tumorigenesis and age-related diseases.

## Introduction

Discovered in cultured human fibroblasts (1), cellular senescence describes a stress response characterized by irreversible cell cycle arrest, accompanied by several major changes in cellular morphology and physiology. Senescence can be induced by a variety of stimuli, such DNA damage, oncogene activation, mitochondrial dysfunction, telomere shortening and oxidative stress (reviewed in (2); (3)).

Apart from cell division arrest, senescent cells (SCs) exhibit a complex phenotype, including cellular hypertrophy, increased ß-gal function (SA-ß-gal) due to enhanced lysosomal activity, resistance to apoptosis, ROS production and heterochromatin modifications. Importantly, they also secrete a large number of factors, including cytokines, chemokines and other small molecules; this property, referred to as Senescence Associated Secretory Phenotype (SASP) (reviewed in (4); (5)), is of special interest, for it is through the SASP components that the SCs communicate and interact with the surrounding tissue.

The SCs are thought to have some beneficial effects; the cell cycle arrest is seen as an antitumor function as it impedes the proliferation of damaged cells after stress. Moreover, SCs are also involved in normal mouse embryogenesis (6) and remodeling and repair (reviewed in (2); (7).

However, the accumulation of SCs has been implicated in some pathologies, like inflammation and cancer (8); (9). Some of the SASP components behave as growth factors and can induce overgrowths, originated by excess of proliferation of non-SCs in the surrounding tissue. In the case of *Drosophila*, the production by SCs of Wg, Dpp and Upd, the ligands of the Wg, Dpp and JAK/STAT pathways respectively, are involved in the generation of tumorous overgrowths (10).

A significant finding is that the number of SCs increase with the age of the organisms and has been conjectured that the accumulation of SCs may be associated with some pathologies associated with old age (11); (12).

Although most studies on senescent cells have been performed in vertebrate tissues or in cells in culture, it has been shown that *Drosophila* epithelial cells, just like mammalian cells, can become senescent in response to mitochondrial defects (13), genetic stress (14), oncogene activation (15), or irradiation (10). Moreover, the *Drosophila* SCs share the same features with vertebrate SCs: cell cycle arrest, cellular hypertrophy, SA-ß-Gal, ROS production, SASP, chromatin modifications, etc (15); (10). This highlights the overall conservation of the process, which makes it possible to use convenient model organisms to study the general properties and behavior of SCs.

Cellular senescence in mammalian cells and in *Drosophila* have many points in common, although there are factors involved in cellular senescence in vertebrates that have not been characterized in *Drosophila.* The transcription factor complex NF-Kß is considered as a determining factor in vertebrates, especially in the acquisition of the SASP phenotype (16); (5). In *Drosophila* NF-Kß has a major role in the innate immune response (reviewed in (17)), but its role in cellular senescence in *Drosophila* has not been examined.

There are two major factors implicated in the acquisition of senescence in *Drosophila*: the Jun N-Terminal Kinase (JNK) pathway and the P53 transcription factor (10). Of these, JNK plays the principal role, for while the lack of *p53* still allows an amount of cellular senescence after IR, no senescence is observed in absence of JNK function. The function of *p53* appears to be subsidiary, propitiating high levels of JNK activity (10).

The activation of JNK by IR-induced DNA damage in *Drosophila* is well known (reviewed in (18)). Double strand breaks are detected by sensors such like the MRN protein complex, which activate the ATM (Ataxia Telangiectasia Mutated) kinase that in turns phosphorylates several substrates, including the kinase Chk2. The latter activates *p53*, which induces JNK expression (19) and the onset of senescence (10).

A remarkable property of SCs, reported in mammalian tissues, is their ability to induce senescence in cells in their proximity, a phenomenon mediated by the SASP (20); (21).

This process of paracrine induction (PI) increases the number of SCs and is likely a contributing factor to the accumulation of SCs with the age of the organisms. In oncogene-induced senescence, Acosta et al (2013) found clear evidence that PI is mediated by multiple components of the SASP, and especially by the Transforming Growth Factor ß (TGF-ß), which activates the CDK inhibitors p15INK4b and p21CLP1, thus blocking cell division. This recruitment mechanism causes accumulation of SCs in the tissues, a factor that may enhance some of the pathologies associated with SCs.

While the phenomenon of PI and mechanisms are well established in mammalian cells (22), its evolutionary conservation has not been tested. We have used the wing imaginal disc of *Drosophila* (which contains the precursors of the wing and thorax of the adult fly) to investigate paracrine recruitment of new SCs. We show that it may account for more than half of the final population of SCs. Moreover, we find that PI requires secondary activation of JNK by a non-canonical role of the Dpp pathway, involving the TGF-beta activated kinase 1 (Tak1), an upstream component that activates the JNK pathway (23). In turn, the activation of Tak1 is achieved by a novel pathway that incorporates several of the components of the immune deficiency pathway (Imd), including the Dredd/Imd/Fadd signaling complex, Tak1 and the NF-kB transcription factor Relish. Our data also suggest that the accumulation of SCs caused by PI is a major factor in senescence-induced tumorigenesis in *Drosophila*.

## RESULTS

### 1) Induction of SCs in the wing disc of *Drosophila*. Role of the JNK pathway

We have recently developed an easy and efficient method to generate SCs in the imaginal discs of *Drosophila* (Figure 1a). It is based on the finding that irradiated imaginal cells that cannot execute the apoptosis program respond by activating the JNK pathway permanently (24), what causes transformation towards senescence of many of those cells (10). In our experiments the cells are made defective in apoptosis by driving the expression of a miRNA (miRHG) construct that inactivates the principal pro-apoptotic genes, *reaper*, *hid* and *grim* (25). The usage of the Gal4/UAS system permits to target a specific area where cells are made apoptosis-deficient.

**Figure 1.**
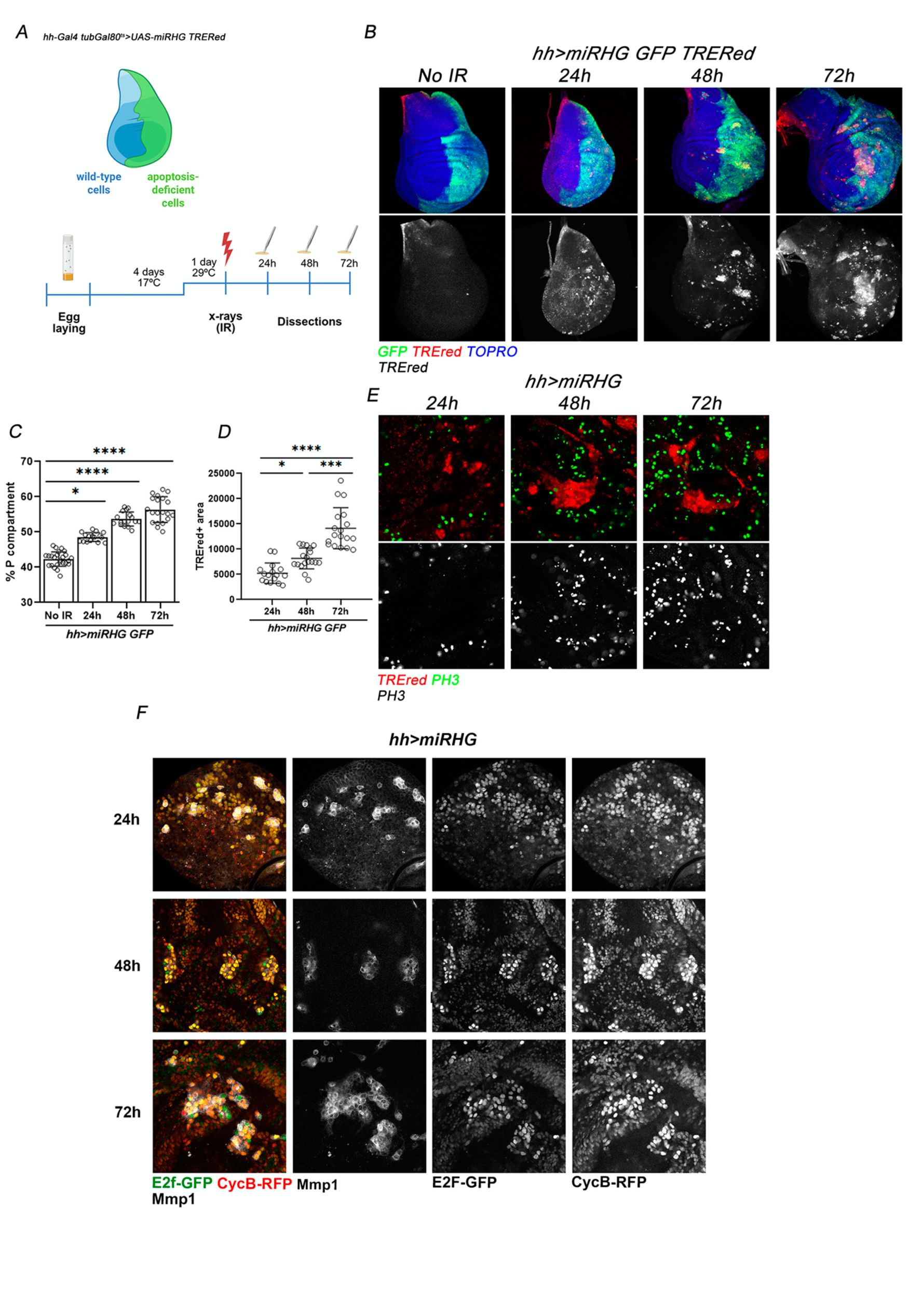
Standard experiment of induction of senescent cells (SCs) by irradiation (IR) A) scheme of the experimental procedure: Irradiated discs of *hh>mirRHG GFP gal80^TS^ TREred* genotype (see Material and Methods for further details) were fixed at different times (24, 48, 72h) after IR. The expression of the miRHG construct in the P compartment cells (labeled green, as driven by the hh-Gal4 line) makes those cells apoptosis-deficient and many of them may become senescent. B) Visualization of SCs, labeled red with the TREred construct, at different times post IR. C) Size increase of the P compartment over time after IR. D) Quantification of the number of SCs at different times post IR. E) Portions of wing discs, 24, 48, 72 h post IR, stained with PH3 (green) as indicator of cells in mitosis. Note that none of the red cells exhibit PH3 staining at any time as quantified in E. F) label with the Flyfucci system to reveal the stage of the cell cycle in with SCs are arrested. SCs are marked here with Mmp1, a JNK target, Ef2 and CycB. Note that at 24, 48 and 72h post IR SCs contain both Ef2 and CycB expression indicating they are arrested at the G2 period shortly after the irradiation.

Using this method, we have followed the appearance in the wing disc of SCs after the irradiation event. The experiment is schematized in Figure 1A, B. Larvae of genotype *hh^Gal80^> miRHG GFP TREred* (see Methods for a full description of the genotype) are irradiated and wing discs are extracted and fixed from larvae at different times after irradiation. Only cells of the Posterior (P) compartment are apoptosis-defective, thus SCs can only appear in the P compartment. Cells of the anterior (A) compartment serve as control. The TREred is an overall marker of JNK activity (26), which we have shown is active in all SCs (10).

The results are illustrated in Figure 1B-D. Because the irradiation causes overall activation of JNK (27); (28), at 24h post IR we find JNK upregulation in both the A and the P compartment, although the staining pattern is different. In the A compartment the TREred label is punctuated and appears in dots of different sizes, suggesting it marks fragments of dead cells; this label disappears after 24h. In contrast, in the P compartment we find TREred label in individual cells, which, as we show below, present signs of senescence. As we reported previously (10), the size of the P compartment gradually increases with respect to the rest of the disc.

Since cell division arrest is the key distinguishing feature of SCs (2); (3), we first used the proliferation marker PH3 to check mitosis in discs at 24, 48 and 72h post IR. The clear result is that JNK-expressing cells do not show PH3 staining at any of those time points (Figure 1E), indicating that by 24h after IR they are already cell-cycle arrested. This is further supported by staining with the Flyfucci system (29). All cells expressing JNK, in this case monitored by the Metalloproteinase 1 (Mmp1) marker, also show expression of E2F and CycB, indicating JNK-expressing cells are arrested at the G2 period (Figure 1F). The lack of PH3 staining and the arrest at G2 appear to affect to all the JNK-expressing cells examined.

In addition, we have examined the appearance of various senescence biomarkers during the 24-72h period post IR. We paid attention to SASP markers already identified in *Drosophila*, like Dpp, Upd and Wg, and to others like cellular hypertrophy or ROS production. The results are illustrated in Figure 2A-F. By 24h post IR, JNK-expressing cells show expression of Upd and Wg, and show ROS production whereas *dpp* expression and cell hypertrophy are detected after 48h.

**Figure 2.**
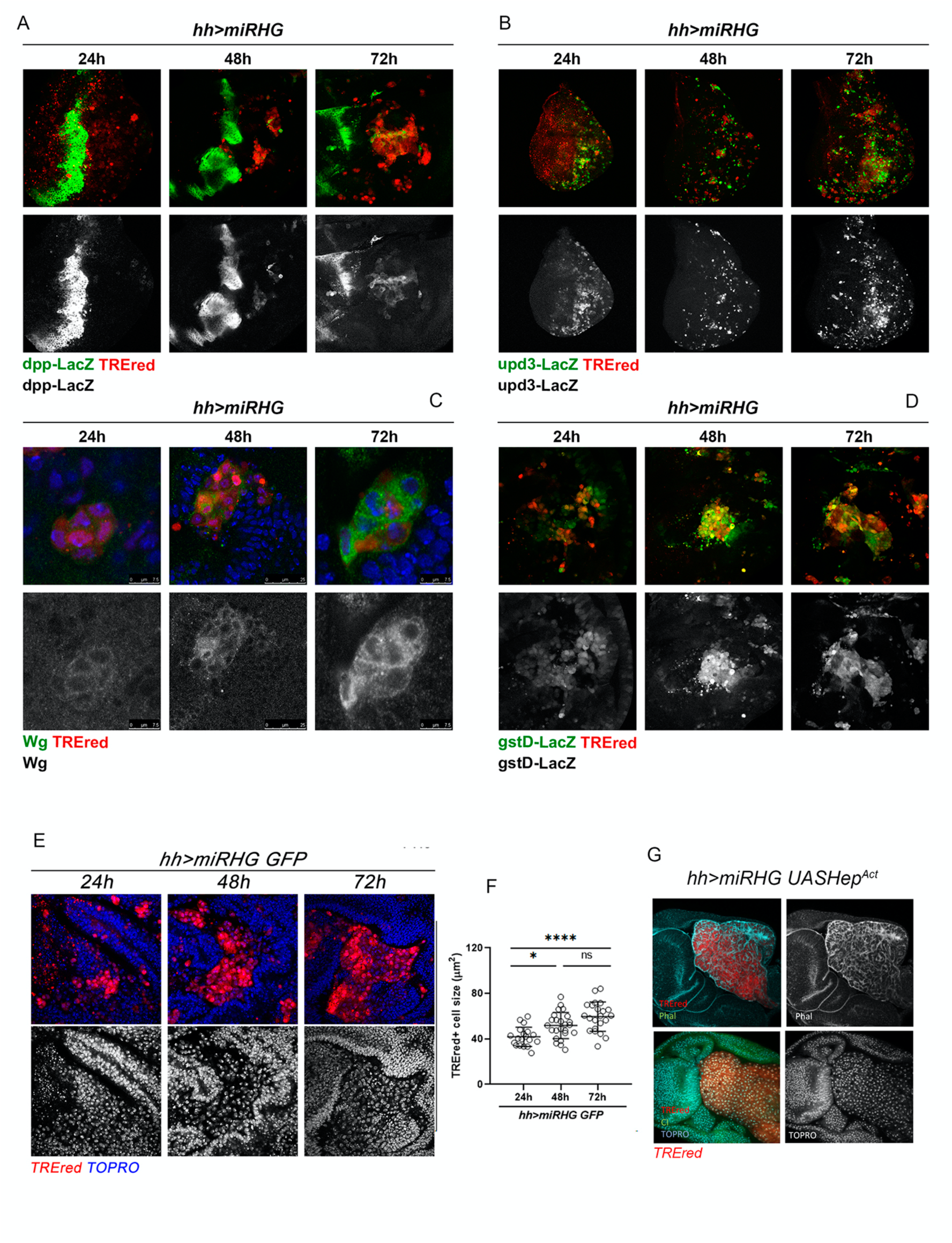
Expression of senescence biomarkers at different times (24, 48 and 72h) after irradiation. Discs of *hh^Gal80^> miRHG TREred* genotype (simplified in the Figure as *hh>miRHG*. In some experiments some additional constructs *(upd-LacZ, dpp-LacZ, gst-LacZ*) have been introduced to detect expression of the corresponding marker. In all figures, JNK-expressing cells are labeled red with the TREred construct. The various biomarkers are labeled green. A) The images show the normal band of *dpp* expression in the middle of the disc and the SCs in red. At 24h post IR none of the red cells show *dpp* expression, but those fixed at 48 and 72h contain *dpp* activity, B) Activity of the *upd* gene is detected in JNK-expressing cells 24h after the irradiation and in subsequent times. Note that all red cells show *upd* expression, C) Expression of the Wg protein is already detected at 24h post IR, although perhaps at lower concentration that at later times. D) TREred cells show ROS production, revealed by the expression of the *gstD* gene, at all the times examined. E) Size increase of TREred cells over time. At 24h post IR there is no detectable difference with non-TREred cells in their proximity but at 48h and especially at 72h, the difference is clear. Quantification in f).

A detailed description of the appearance of senescence biomarkers during the transition towards senescence is pending, but the significant observation is that 24h after the irradiation, JNK-expressing cells exhibit activity of key markers, clearly indicating that they have attained senescence status shortly after IR. We also note that all the JNK-expressing cells we have examined show senescence markers, indicating that JNK expression is a reliable marker of cellular senescence in *Drosophila*.

Because in all the preceding experiments the JNK pathway appeared to be associated with the senescence phenotype, we wondered whether its sole activity would be sufficient to induce senescence in apoptosis-deficient cells. We tested this in an experiment in which we forced JNK activity in the P compartment, using the hh>Gal4 line. To this effect we used a constitutive form of the Hemipterus kinase *hep^Act^*, a key activator of the JNK pathway (30). The result is shown in Figure 2G: in *hh> hep^Act^ miRHG TREred* discs virtually all the cells of the compartment become senescent, as judged by their morphology and size. This is a significant result, for it assigns to JNK a principal role in the establishment of cellular senescence in *Drosophila*.

There are two main conclusions from these experiments, 1) JNK-expressing cells acquire senescence status shortly after the triggering event, although full implementation may take some more time, 2) JNK is a major inducer of cellular senescence in *Drosophila* for its sole activity is sufficient to trigger senescence. This conclusion is especially relevant, for in subsequent experiments we count JNK expression as a general marker of senescence.

### 2) Increase of SCs by paracrine recruitment mediated by Dpp

In addition to the size increase of the P compartment of irradiated *hh^Gal80^> miRHG GFP* TREred discs (10), we find that the number of JNK-expressing cells in the compartment also increases over time. We have estimated their number indirectly by measuring the surface covered by TREred tissue in the compartment and comparing it with the non-TREred area. The comparison was made in discs fixed at 24, 48, 72 after IR.

Initially (24h post IR), the SCs area covers approximately 10% of the compartment, but it gradually rises with time to reach a maximum of 24% in the 72h series (Figure 3A-B). Besides, because SCs do not divide (Figure 1,E,F), this result indicates that the increase of the SCs population is due to recruitment of new SCs, most likely by induction by pre-extant SCs.

**Figure 3.**
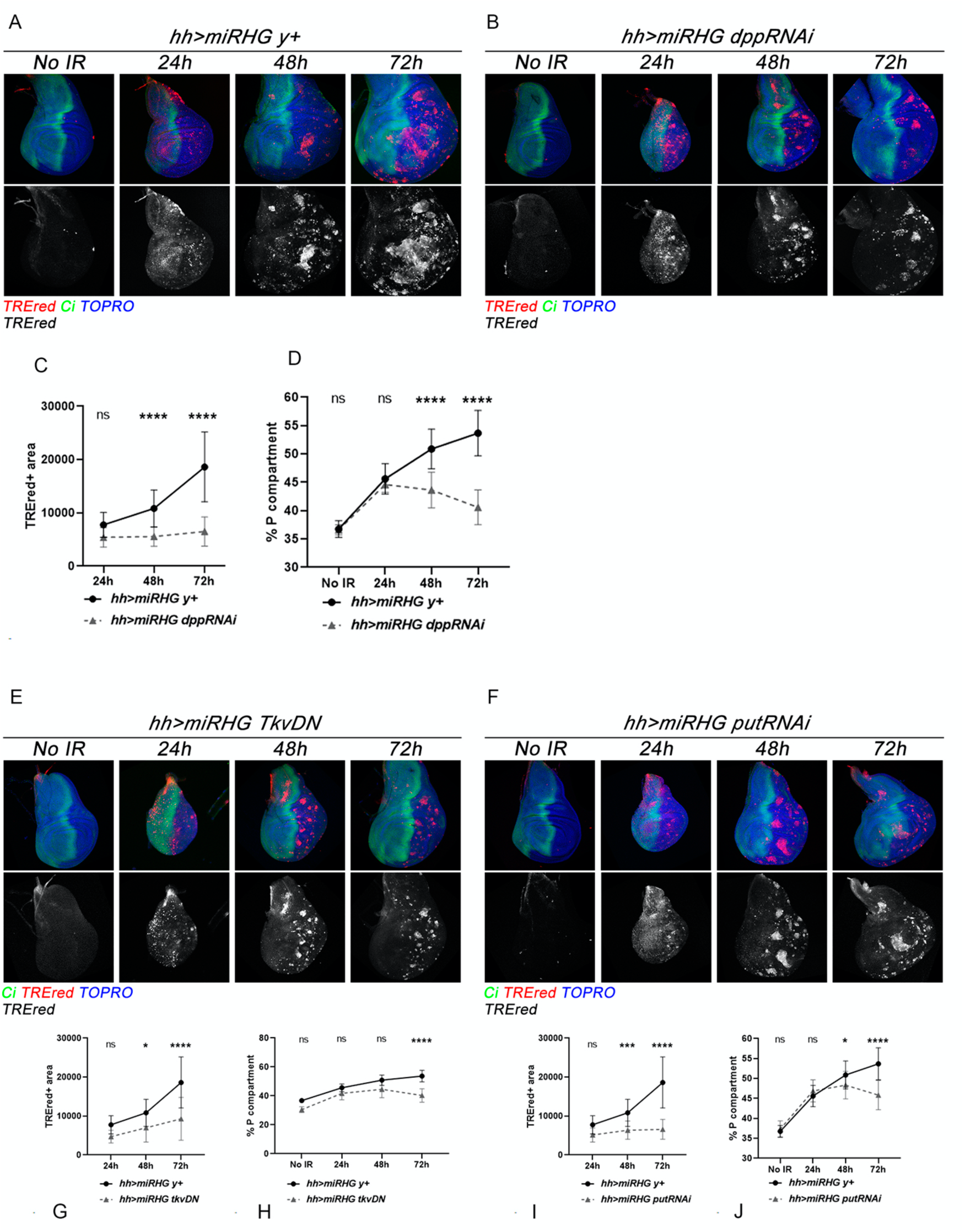
Effect of the suppression of the Dpp pathway on the emergence of new SCs A) Control experiment. Discs of genotype *hh^Gal80^> miRHG GFP TREred* (genotypes in the figures are simplified for clarity) show the normal increase of SCs in the P compartment over time after IR (also illustrated in Figure 1B). The A/P border is delineated by the expression of the *cubitus interruptus (ci)* gene in green. B) Discs of similar genotype in which the *dpp* function is suppressed by an effective RNAi line. Note that the number of SCs remain constant after 24h, as quantified in the C panel. D) The lack of *dpp* activity in SCs impedes the overgrowth of the P compartment. Compare with the control *hh>miRHG y^+^.* E) Consequences of suppressing the function of the Tkv receptor. In *hh^Gal80^> miRHG GFP TREred Tkv^DN^* discs there is only a small increase of SCs with time, much less than in the control *hh^Gal80^> miRHG GFP TREred* y*^+^* and the growth of the P compartment is also reduced in comparison with the control (quantifications in the bottom row), F) The lack of function of the receptor Punt also prevents the accumulation of SCs over time and the overgrowth of the P compartment is much reduced. G-J) Quantification of the results of the receptors experiments

We surmised that the increment of SCs over time could be due to some SASP component emanating from the initial cohort of SCs. In mammalian cells there is evidence of the implication of SASP components in PI, with especial relevance of the TGF-ß secreted factor (20). We then focused on Dpp, the *Drosophila* ortholog of the mammalian TGF-ß.

Using RNAi constructs that compromise the production of Dpp by the SCs, we have assayed whether it is necessary for the induction of new SCs. In those experiments we have estimated the number of SCs at 24, 48 and 72h after IR. The evolution of the SCs population after IR in *hh^Gal80^>miRHG TREred dpp^RNAi^* discs is shown in Figure 3B,C; the significant finding is that the number of SCs remains at the 24h level and does not increase over time. We examined those cells to certify that they are indeed SCs; they do not express cell proliferation markers like PH3 and show SA-ß-gal activity. The conclusion is that in the absence of the Dpp signal JNK-expressing cells can become fully senescent, but are unable to amplify their number.

These results indicated that there is a first wave of SCs generated by irradiation-induced JNK activity, followed by a second wave, 48h onwards, mediated by the Dpp signal. Since JNK activity is a principal inducer of senescence, these observations in turn suggested that Dpp is involved in a secondary activation of JNK that raises the number of SCs.

Additional experiments tested the implication of the Dpp pathway; we compromised the transduction of the pathway in the P compartment by blocking the activity of the Thick vein (Tkv, type 1) and Punt (Put, type 2) receptors (reviewed in (31)) using a dominant negative form of Tkv and a putRNAi line. In discs of *genotype hh^Gal80^>miRHG TREred UAS-tkv^DN^* or *hh^Gal80^>miRHG TREred UAS-putRNAI* we find no increase of SCs over time (Figure 3E-J); their number remain at 24h level for the rest of development, essentially the same result found when the Dpp ligand is absent. This finding indicates that the activation of the Dpp pathway is a necessary step for non-SCs cells to undergo senescence.

The experiments described in this section demonstrate two distinct requirements of Dpp functions in the process of paracrine induction: 1) Pre-extant SCs have to be able to generate and secrete the Dpp signal. This has already been shown (10) and is a feature of JNK-expressing cells (Pinal et al 2019), and 2) the Dpp pathway has to be activated in receiving non-SCs to undergo senescence, likely by activating JNK.

### 3) A non-canonical role of Dpp induces secondary activity of JNK. The Imd route

The experiments above call for a role of the Dpp pathway in the secondary JNK activation that generates new SCs. Hence, we searched for mechanisms by which Dpp may regulate JNK.

There is evidence that Dpp controls ommatidia differentiation by a non-canonical route that results in the activation of the Transforming growth factor Activating Kinase 1 (Tak1), which in turn induces activity of the JNK pathway, followed by upregulation of the Matrix metalloproteinase 1 (Mmp1). This cascade of events occurs without the intervention of the Mad/Medea effector (32).

Consequently, we explored the possible implication of the Tak1 kinase in paracrine induction of SCs. First, we checked whether Tak1 is able by itself to induce JNK activity. The result is illustrated in Figure 4A: driving Tak1 expression in the P compartment of non-irradiated *hh>miRHG TREred Tak1* discs results in high level activity of JNK, as indicated by the TREred label. Thus, the sole activation of Tak1 in the P compartment in *hh>miRHG TREred Tak1 d*iscs causes the same syndrome as the irradiation of *hh>miRHG TREred* discs. Since Tak1 is an upstream element of JNK, this finding reinforces the notion that JNK is a primary inducer of cellular senescence in *Drosophila*.

**Figure 4.**
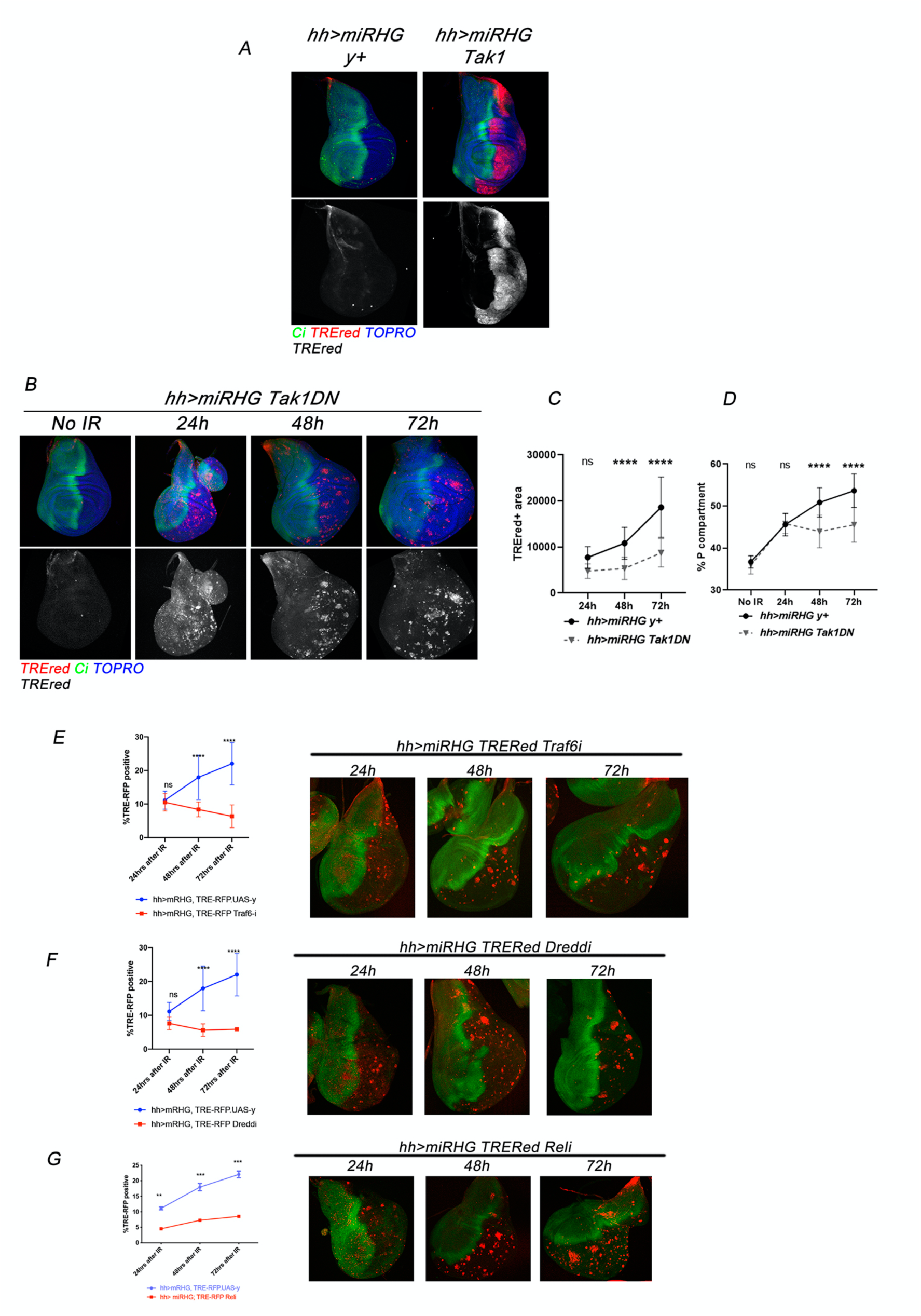
Role of Tak1 and components of the immune response pathways in paracrine induction of SCs. A) Driving Tak1 expression in the P compartment in discs of genotype *hh^Gal80^> miRHG GFP TREred UAS-Tak1* causes high levels of JNK activity in most of the compartment cells. The A/P line is delineated by the expression of *ci*, a marker of the A compartment (green), B) Loss of TAk1 activity in the P compartment by forcing expression of the Tak1DN construct. The number of SCs remains essentially constant during the 24-72h post IR period, as quantified in the C panel. D) There is no size increase in the P compartment over time. E) Effect of the lack of Traf6 function on the amount of SCs (in red) over time. Unlike in the control (blue line), in the experimental series (red line) there is no increase in the number of SCs, F) Results of a similar experiment removing Dredd function. There is no increase of SCs, G) The lack of Relish function also impedes the increase of SCs with time after IR

Second, we tested whether paracrine recruitment of new SCs requires Tak1 function. We generated SCs in P compartments in which the activity of the Tak1 kinase is compromised by either a dominant negative form or by an effective RNAi line. In both cases we obtain the same result (Figure 4B-C): the number of SCs present at 24h post IR does not increase over the 24-72h period, clearly indicating lack of PI. The overgrowth of the P compartment is also much reduced (Figure 4D)

The results with Tak1 led us to explore the mechanisms of Tak1 activation in the context of paracrine induction. The Tak1 protein is a component of the Immune Deficiency (Imd) pathway of *Drosophila*, which responds to infection of Gram-negative bacteria (reviewed in (17). Imd is triggered by the proteoglycan (PGN), present in most bacteria, which is recognized by the PGN Recognition Proteins (PGRPs). PGN binding to the PGRP receptor causes the recruitment of a signaling complex that includes the adaptor proteins Imd and Fadd, and the caspase Dredd. Cleavage by Dredd allows ubiquitination of Imd (see detailed description of the cascade in (17)). This results in the recruitment of the Tab2/Tak1 complex, which eventually leads to the activation of the NF-kB protein, Relish, which induces function of Anti-Microbial Peptides (AMP), essential for the immune response. In addition to its antimicrobial role, the Tak1 protein is also involved in remodeling *Drosophila* airways by upregulating the JNK pathway through the activation of the upstream kinase Hep (33).

We entertained the hypothesis that some elements of the Imd or of other Immune Response pathways activated by Dpp could be involved in the secondary JNK activation. Consequently, by utilizing RNAi constructs that suppress or weaken their function, we have analyzed the requirements for PI of some key components of the Immune Response pathways. We tested the TNF-receptor-associated factor Traf6 that in vertebrates transduces Toll-receptor signaling (34) and promotes JNK activation (35), the caspase Dredd (part of the Imd/Fadd/Dredd signaling complex of the Imd pathway (17), and the NF-Kß transcription factor Relish. The result is that after compromising the activity of any of the three proteins, PI is impeded (Figure 4E-G) and no JNK activity observed. These observations indicate that in *Drosophila* the PI process incorporates the core of the Imd pathway, including the postransductional modifications of the Imd protein and the activation of the Tak1 kinase, which in turn activates JNK and the NF-Kß Relish transcription factor (see Figure 4G). The finding that Relish is involved in PI is significant, for it demonstrates the implication of NF-kB in the SASP phenotype in *Drosophila*, just as in mammalian cells (36). Another indication of the general conservation of cellular senescence in metazoans.

Altogether, the functional assays of the different components of the innate immune system suggest that several of them have been co-opted in *Drosophila* to build a specific pathway to ensure Dpp-dependent activation of JNK, necessary to generate a secondary wave of SCs (see Discussion).

The implication of a non-canonical function of Dpp in paracrine induction of new SCs is intriguing because we find that the canonical function is also active. First, the pMad effector (31) is activated at low level in SCs, but more strongly in non-SCs in the vicinity (Figure 5A). The target gene *daughters against dpp* (*dad*) is expressed at high levels in all SCs (Figure 4B). Another Dpp target, *brinker (brk*) is expressed in SCs and also in non-SCs neighbors (Figure 5C). The results with pMad and *brk* indicate paracrine induction mediated by the canonical route of Dpp.

**Figure 5.**
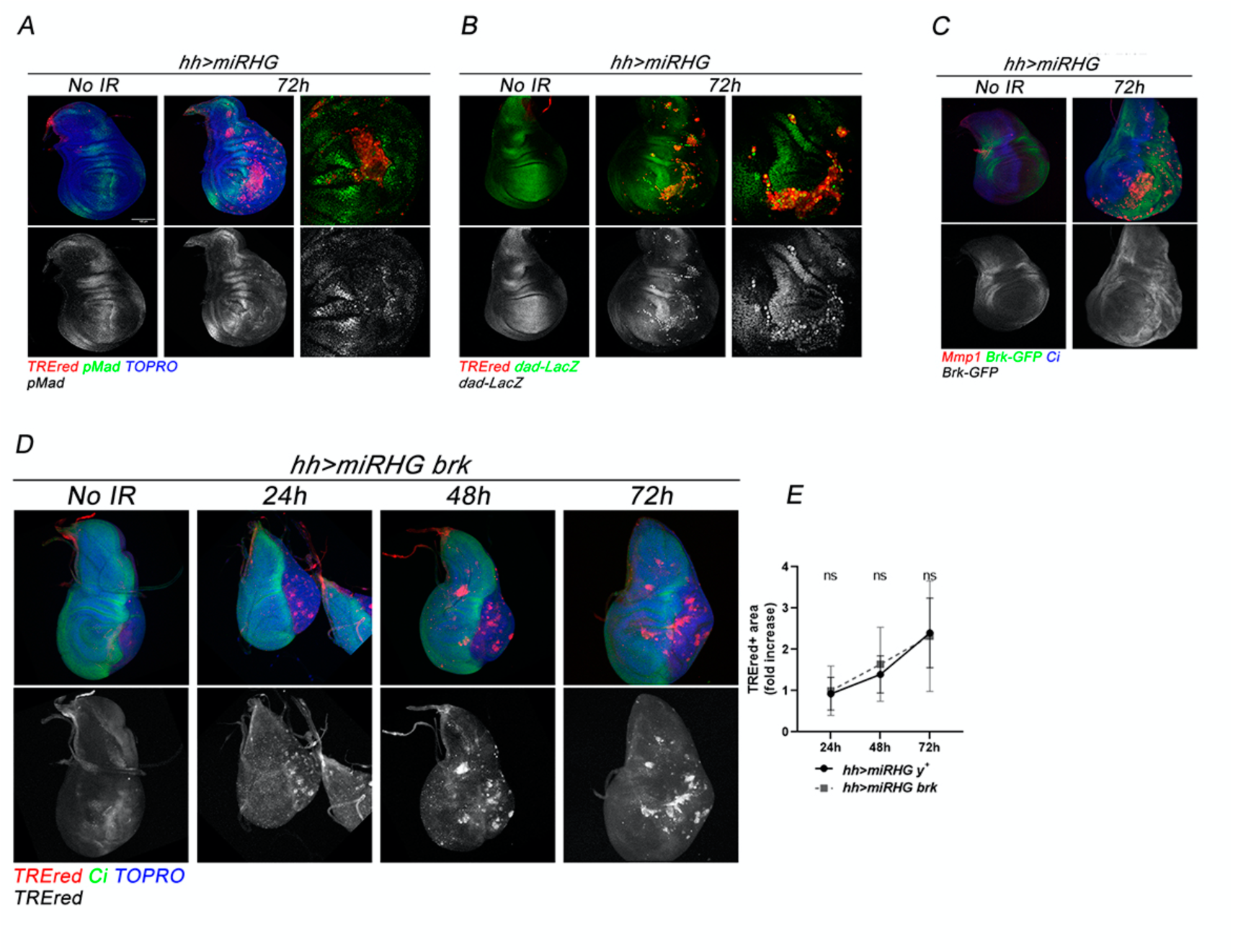
The Dpp canonical route in irradiated *hh>miRHG TREred* discs 72h after the irradiation. Expression of various components of the route. A) the pMad effector (green) is expressed at low levels in the SCs, but at higher levels in neighbors non-SCs, indicating that the latter have triggered the Dpp pathway. The image to the right is a high magnification to illustrate the point more clearly. B) the target gene *daughters against dpp* (*dad)* is expressed at high levels in the great majority of the SCs, whether they are inside the normal Dad domain or outside. This image indicates that at least some components of the Dpp canonical are active in SCs, C) The target gene *brinker (brk)* is expressed in SCs at low levels and more robustly in non-SCs in the vicinity. D) The overexpression of *brk* in the P compartment does not prevent the rise of SCs over time after IR. The amount of TREred tissue is not significantly different from control, illustrated in the graph in panel E.

Thus, it seems that both the canonical and non-canonical functions of the Dpp pathway are active in SCs and non-SCs. We have tried to discriminate between those two Dpp functions, by overexpressing the negative regulator *brk*, which effectively blocks the canonical route, mediated by Mad/Medea (37). In irradiated *hh^Gal80^>miRHG TREred UAS-brk* discs the number of SCs increases with time after the IR (Figure 5D,E indicating that there is paracrine induction.

These results reveal two distinct roles of Dpp signaling in tissues containing SCs. 1) Paracrine induction of new SCs, which depends on the non-canonical route and operates through a cascade of events that incorporates components of the Immune Response pathways. This leads to secondary activation of JNK and recruitment of additional SCs. 2) The canonical Dpp route, in contrast, is involved in growth and is likely responsible for the formation of tumorous overgrowths generated by SCs (9)

## Discussion

### 1) The JNK pathway as senescence inducer in Drosophila

Let to consider the features of our system to generate senescent cells in *Drosophila* (10). It is based on provoking a physiological stress, generally irradiation (Figure 1A), in cells that cannot execute the apoptosis program. The fact that cells are apoptosis-deficient is essential because IR activates JNK, what provokes cell death by apoptosis. The suppression of the apoptosis program allows the survival of irradiated cells. Besides, the cells with high levels of JNK activity, which presumably would have been killed by IR, retain JNK function for the rest of the development, what causes tumorous overgrowths (24). The persistence of JNK activity is due to the establishment of a JNK maintenance loop by *moladietz (mol)* and ROS (24)(38).

Our experiments demonstrate the key role of the JNK pathway triggering cellular senescence in *Drosophila*. First, the cells that contain JNK activity also show senescence features (Figures 1 and 2). Second, and importantly, just driving the constitutive form of the Hemipterous kinase Hep^Act^, a key activator of the JNK pathway, is sufficient to trigger the transformation towards senescence (Figure 2G). Our experiments also show that in our experimental system JNK-expressing cells acquire senescence features, cell cycle arrest, ROS production, Wg and Upd activity, shortly (24h) after the triggering event, while other senescence biomarkers may appear later.

### 2) Increase of SCs over time after the initiation event. Paracrine induction of SCs by a non-canonical route of Dpp

A key result is that the number of SCs increases during development after the stress event: 24h after IR the SCs cover 10% of the P compartment of *hh^Gal80^> miRHG GFP TREred* discs, while by 72h they cover 24%. Since the SCs are cell-cycle arrested, it follows that the increase in cell numbers must occur through a mechanism other than cell division. This phenomenon cannot be attributed to a delayed effect of irradiation, since the effects of X-rays dissipate within 24–48 hours. Besides, the fact that in absence of *dpp* function the number of SCs remains at the 24h level for the whole 24-72 period argues strongly against the latter possibility.

Our results establish the role of the Dpp pathway in the recruitment of SCs by paracrine induction. This is in line with previous work in mammalian cells (19), which demonstrated the implication of TGF-ß (the vertebrate ortholog of Dpp) in the same process. If the emitting SCs cannot secrete the Dpp signal or the receiving non-SCs cannot activate the Dpp pathway (for example, lack of function of the Tkv/Put receptors), the induction of new SCs is prevented (Figure 3 C,G,I). The lack of Dpp function does not prevent cells from become senescent, but impedes the recruitment of new SCs.

These findings allow distinguishing two waves of SC (Figure 6): 1) First-generation SCs, in which JNK activation is directly induced by irradiation, 2) Second-generation SCs, in which JNK is activated later in non-SCs through Dpp signaling. This is achieved by a non-canonical route of the Dpp pathway, which requires the Tkv/Put receptors, but branches off to activate the Tak1 kinase, which in turn activates JNK and the NF-Kß transcription factor Relish (Figure 6). It is noteworthy that the cascade of molecular events leading to the activation of Tak1 and Relish during the PI process mirrors that of the Imd signaling pathway, which mediates the immune response to Gram-negative bacterial infection, although it incorporates factors like Traf6 that in vertebrates is part of the Toll pathway. This observation exemplifies how evolutionarily conserved signaling modules can be co-opted and repurposed to fulfill distinct biological functions. Although there is the possibility that Traf6 and Imd may activate JNK independently, our finding that there is no paracrine induction when either of them is lacking suggests that in *Drosophila* they are interdependent e.g. they may be part of the same cascade.

**Figure 6.**
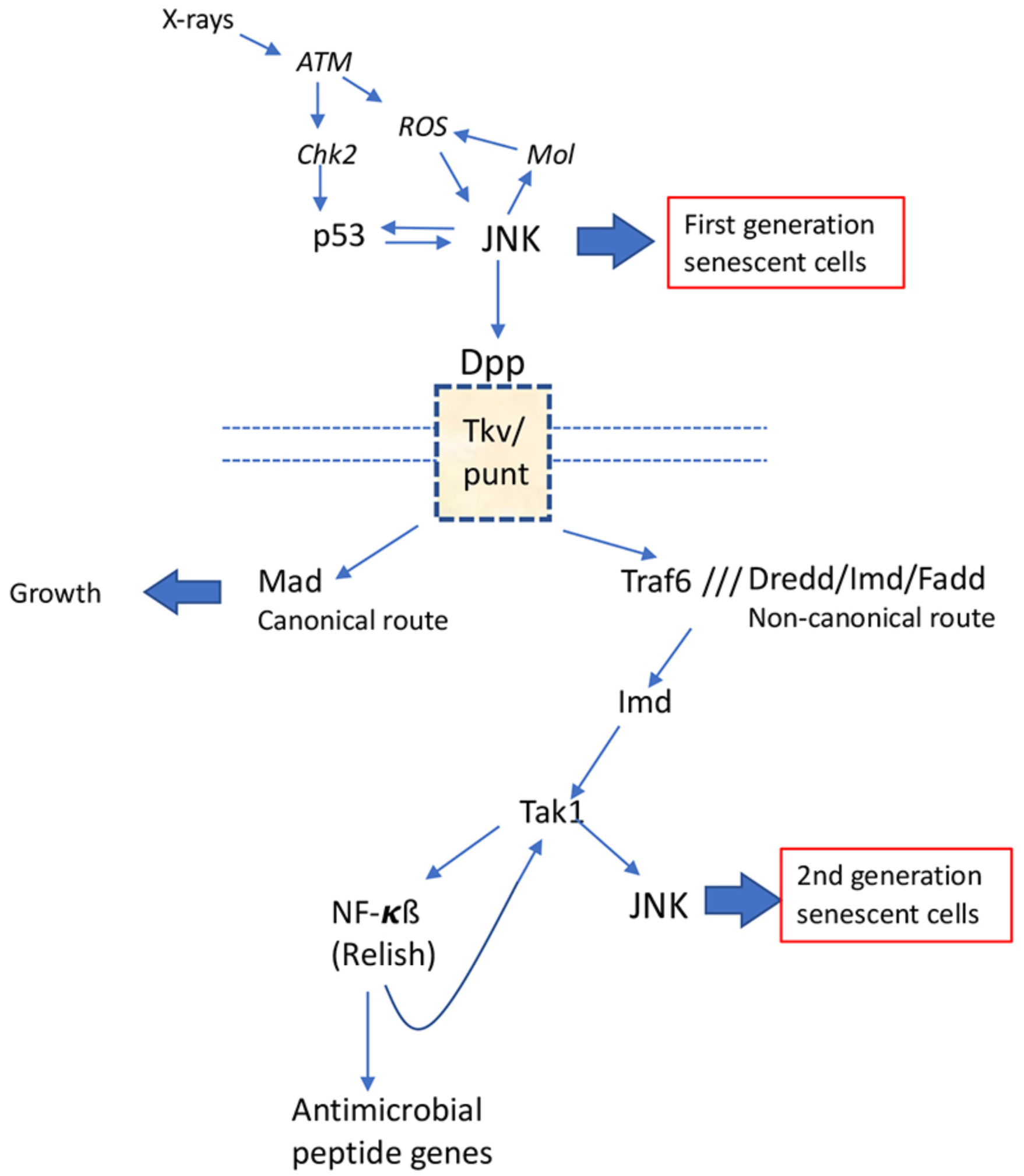
JNK as a principal inducer of SCs *Drosophila*. Irradiation causes double strand break that are detected by sensors such like the MRN protein complex, which activate the ATM (Ataxia Telangiectasia Mutated) kinase that in turns phosphorylates several substrates, including the kinase Chk2. The latter activates *p53*, which induces JNK expression and the onset of senescence in a group of imaginal cells, the first generation of SCs. Those SCs release a SASP component, the Decapentaplegic (Dpp) signal, which is transduced through the Tkv/Punt receptor complex and activates the Dpp pathway in non-SCs in the proximity. For unknown reasons some non-SCs choose the Mad-mediated Dpp canonical route, which causes tissue growth. Other cells take a non-canonical route involving the kinase Tak1, which induces JNK activity and originates a second wave of SCs. The activation of Tak1 is achieved by a novel pathway that incorporates several of the components of immune deficiency pathways including the Dredd/Imd/Fadd signaling complex and the NF-kB transcription factor Relish.

The fact that the NF-Kß factor Relish is necessary for paracrine recruitment fits well with the evidence from the vertebrate field that NF-Kß plays a major role in the induction of the SAS Phenotype (39). Based on our finding that lack of Relish function prevents paracrine induction, we propose the existence of a regulatory loop by which Relish stimulates Tak1 activity.

Although our results clearly demonstrate the requirement of Dpp for paracrine induction of SCs, they do not exclude the contribution of other SASP components, as is the case in mammalian cells (20).

#### Dual role of Dpp in cellular senescence

Finally, there is the observation that in *Drosophila* the Dpp signal can be transduced differentially by non-SCs of the same tissue, with different consequences. Most cells proceed through the canonical route, remain non-SC and are able to proliferate. But a smaller subset instead follows the non-canonical route, undergoing Tak1- and JNK-dependent induction of senescence. This differential function has two distinct consequences, illustrated in Figure 9: 1) through the canonical route, Dpp functions as a growth factor, stimulating cell proliferation of non-SCs, presumably by the SCs secreting proliferative signals, including Dpp itself. This causes the formation of tumorous overgrowths when there is a sufficient number of SCs. The role of Dpp as a growth factor in the wing disc is well established in the literature (Affolter and Basler 2007). 2) through the non-canonical route, Dpp amplifies at least twofold the original number of SCs generated by the initial inducing event. At this stage we do not know how these alternatives are chosen.

These two roles are interconnected regarding the tumorigenesis generated by SCs. The canonical route generates the signals that stimulate the proliferation of non-SCs, but the non-canonical route increases the number of SCs, thus enhancing the production of proliferative signals. If paracrine induction is prevented the number of SCs is not sufficient to induce tumorigenesis

## Methods

### Drosophila strains

All the *Drosophila* strains employed in this study were maintained on standard medium at 25 °C (except for the temperature shift experiments, see below). The following strains were used: *hh-Gal4* (Tanimoto, 2000), *tub-Gal80^ts^* (McGuire, 2003), *UAS-miRHG* (Siegrist, 2010), *UAS-y^+^* (Calleja, 1996), *TREred* (Chatterjee, 2012), *UAS-GFP* (BDSC#4775), u*bi-GFP-E2f1^1-230^, ubi-mRFP1-NLS-CycB^1-266^* (Fly-Fucci, BSDC#55099), *gstD-LacZ* (Sykiotis, 2008), *dpp-LacZ* (p10638, BDSC#12379), *UAS*-*dpp^RNAi^* (Haley, 2008), *Upd3-LacZ (*BSDC#98418), *UAS-hepAct* (BSDC#*9306)*, *UAS-tkv^DN^* (Vienna stock center #3059), *UAS-put-RNAi* (BSDC#39025), *UAS-Tak1-HA* (BSDC#*59003), UAS-Tak1^DN^ (*BSDC#*58811), UAS-Tak1-RNAi (*BSDC#31045), *UAS-omb-LacZ (*BSDC#*83158), UAS-dad-Lac (*BSDC#*10305), Brk-GFP (*BSDC#*38629), UAS-Brk (*BSDC#*93081), UAS-Imd-RNAi (*BSDC#38933), *UASDredd*-*RNA*i (BSDC#99885), *UASFadd-RNAi* (BSDC#55099), *UASRel-RNAi* (BSDC#33661), *UASTraf6-RNAi* (BSDC#33931).

### Imaginal discs staining

Third instar larvae were dissected in cold PBS and fixed with a mix of 4% paraformaldehyde, 0.1% deoxycholate (DOC) and 0.3% Triton X-100 in PBS for 27 min at room temperature. After that, the samples were washed 4 times in PBS, 0.3% Triton X-100 and blocked in PBS, 1% BSA, and 0.3% Triton X-100 and incubated with the corresponding primary antibody at 4 °C over night. The they were washed 4 times in PBS, 0.3% Triton X-100 and incubated with the corresponding secondary antibodies for 2 h at room temperature in the dark. They were then washed 4 times in PBS, 0.3% Triton X-100 in the dark and mounted in Vectashield mounting medium (Vector Laboratories).

The following primary antibodies were used: rat anti-Ci (DSHB 2A1) 1:50; rabbit anti-PH3 (Cell Signal Technology) 1:100; mouse anti-Mmp1 (DSHB, a combination, 1:1:1, of 3B8D12, 3A6B4 and 5H7B11) 1:50; mouse anti-Wingless (DSHB 4D4) 1:50; mouse anti-b-galactosidase (DSHB 40-1a) 1:50; rabbit anti-pMad (1:100).

Fluorescently labelled secondary antibodies (Molecular Probes Alexa-488, Alexa-555, Alexa-647, ThermoFisher Scientific) were used in a 1:200 dilution. Phalloidin TRITC (Sigma Aldrich) and Phalloidin-Alexa-647 (ThermoFisher Scientific) were used in a 1:200 dilution to label the actin cytoskeleton. TO-PRO3 (Invitrogen) and DAPI (MERCK) was used in a 1:500 dilution to label the nuclei.

### b-galactosidase activity assay

To detect senescence-associated ß-gal activity (SA- ß-gal activity), a CellEvent^TM^ Senescence Green Detection Kit (Invitrogen) was used. After dissection, larvae were fixed in 4% paraformaldehyde for 20 min and, after washing with PBS, 1% BSA, incubated with the Green Probe (1:1000 dilution) for 2 h at 37°C in the dark. Then, samples were washed with PBS and then wing imaginal discs were mounted in Vectashield. Images were taken shortly after the protocol to avoid loss of fluorescence.

### Generation of senescent cells in the wing imaginal disc

After a 24-hour egg laying (EL), *Drosophila* larvae of a standard genotype *TREred, UAS-miRHG; hh-Gal4, tub-Gal80^ts^* were kept at 17°C for 3-4 days to avoid activation of the Gal4-UAS system during early larval stages. Then, they were transferred to a restrictive temperature of 29 °C for 24 hours to allow for the activation of the Gal4-UAS system before being irradiated. After that, they were maintained at 29ªC and wandering third-instar larvae were collected for wing imaginal disc dissection at 24, 48 and 72 hours after irradiation. For characterizing the temporal evolution of senescent cells, a *UAS-GFP* construct was co-expressed with the standard genotype to label the posterior compartment. For analyzing the contribution of distinct genes to the process of paracrine induction, a *UAS-y+* construct co-expressed with the standard genotype was used as a control to titrate the number of UAS.

### IR treatment

Larvae were irradiated using an X-ray machine Phillips MG102 at a standard dose of 4000Rads (R), at the developmental time previously indicated (see “Generation of senescent cells in the wing imaginal disc” section)

### Image acquisition, quantifications and statistical analysis

After wing imaginal disc staining, images were captured with a Leica (Solms, Germany) LSM510 vertical confocal microscope and a Nikon A1R inverted confocal microscope. Tissue sections obtained were of an average of 1,5μm width. Image quantification and processing were performed using the Fiji/ImageJ (https://fji.sc) and Adobe Photoshop software, respectively. Schemes in Fig.1 were made with BioRender(https://www.biorender.com/).

To measure the size of the posterior compartment (named as “%P compartment”), a Z-maximal intensity projection was made for each image. Then, the area of the compartment (labelled with GFP or the absence of Ci staining, depending on the experiment) was measured by using the “Area” tool and normalized dividing between the total disc area (labelled by TOPRO staining). For the quantification of JNK activity levels (named as “%TREred*”*), *TREred* positive area in the posterior compartment was first defined with the “Threshold” tool and then measured as the area fraction in the compartment.

Cell size was measured as the average area of a minimum of 10 *TREred* positive cells (labelled by phalloidin staining) and the mean of the area of 10 *TREred* negative cells per imaginal disc.

Statistical analysis was performed using the GraphPad Prism software (https://www.graphpad.com). To compare between two groups, a non-parametric Student’s *t*-test test was used. To compare between more than two groups, a non-parametric, a two-way ANOVA test was used. Sample size was indicated in each figure legend.

## Notes

### Competing Interest Statement

The authors have declared no competing interest.

### Summary of Updates

We have modified the size of the figures and have added the figure legends

### Funding

- Fundacion Ramon Areces ref CIVP225518725

